# Megafauna diversity and functional declines in Europe from the Last Interglacial (LIG) to the present

**DOI:** 10.1101/2023.06.12.544580

**Authors:** Marco Davoli, Sophie Monsarrat, Rasmus Pedersen, Paolo Scussolini, Dirk Nikolaus Karger, Signe Normand, Jens-Christian Svenning

**Affiliations:** Center for Ecological Dynamics in a Novel Biosphere (ECONOVO) & Center for Biodiversity Dynamics in a Changing World (BIOCHANGE), Department of Biology, Aarhus University, 8000 Aarhus C, Denmark; Institute for Environmental Studies, Vrije Universiteit Amsterdam, De Boelelaan 1111, 1081 HV Amsterdam, Netherlands; Swiss Federal Institute for Forest, Snow and Landscape Research WSL, Zürcherstrasse 111, 8903 Birmensdorf, Switzerland

## Abstract

**Aim:** to estimate European megafauna diversity and functional declines in the present compared to the nearest in time modern-analogue climate period prior to the worldwide diffusion of *Homo sapiens*.

**Location:** Europe.

**Time period:** Last Interglacial (LIG; ca. 127,000 years ago) to present.

**Major taxa studied:** wild, large (≥10 kg) terrestrial mammals.

**Methods:** we assessed the distribution of 48 European megafauna species during the LIG using hindcasting modelling and fossil records. Then, we estimated the decline in megafauna community diversity and potential trait-based functional effects from the LIG to the present, accounting for climate differences between the two periods.

**Results:** from the LIG to the present, species richness and community biomass across Europe dropped by 74.3% (± 9.9% SD) and 96.7% (± 4.1% SD), respectively. Functional diversity dropped by 59.1% (± 11.8% SD) for herbivores and by 48.2% (± 25.0% SD) for carnivores, while trait-informed potential vegetation and meat consumptions dropped by 91.1% (± 7.4% SD) and 61.2% (± 17.2% SD), respectively. The loss in megafauna diversity and associated ecological processes were high everywhere, but particularly in western Europe for carnivores and in the East European Plain for herbivores. We found that potential megafauna richness and functional patterns in the two periods were near identical if only climate-driven differences were considered.

**Main conclusions:** severe, size-biased defaunation has degraded megafauna assemblages and megafauna-mediated ecological processes across Europe from the LIG to the present. These patterns cannot be explained by climate differences between the two periods, thus were likely driven by the impact of prehistoric Homo sapiens. The results suggest that the structure of wild ecosystems of the present strongly deviates from the evolutionary norm, notably with decreased functional heterogeneity and decreased fluxes of biogeochemical compounds across the trophic networks, highlighting the importance of ambitious policies of megafauna community restoration to support ecosystems functioning.

## 1. Introduction

Megafauna are disproportionally important for the functioning of biological communities due to a broad range of effects linked to their large body size (Malhi *et al*., 2016; Enquist *et al*., 2020). They influence vegetation structure and dynamics (Bakker *et al*., 2016; Pringle *et al*., 2016), plant migration (Fricke *et al*., 2022b), species diversity (Ratajczak *et al*., 2022), fire regime (Karp *et al*., 2021), nutrient fluxes (Doughty *et al*., 2016), and long-term carbon storage (Kristensen *et al*., 2021). Functionally diverse megafauna communities were once prevalent globally, yet they have been severely downgraded due to worldwide extinctions in the late-Quaternary (Smith *et al*., 2019). Although different causes of these extinctions have been long debated (Koch & Barnosky, 2006), a broad range of evidences increasingly attest a prominent role of the impact of *Homo sapiens* spreading out of Africa (Sandom *et al*., 2014a; Smith *et al*., 2019; Andermann *et al*., 2020). As the result of these extinctions, megafauna worldwide is strongly diminished in numbers, with smaller species remaining, and overall simplified communities relative to the norm of the last millions of years (Smith *et al*., 2018).

While the macroscale patterns of the late-Quaternary megafauna losses are increasingly clear, the pre-extinction megafauna distributions and the drop in the present of associated ecological effects are still incompletely understood. A range of studies provide insights about megafauna role on past ecosystem structure and functioning at local scale, e.g., large-herbivore heterogeneity and vegetation structure in Britain (Sandom *et al*., 2014b), or large-herbivore abundance, fire regime and vegetation composition in Queensland (Rule *et al*., 2012). A global-scale study suggests a general rise in fire activity subsequent to megafauna declines (Karp *et al*., 2021). However, the literature remains limited, in large part because the paleoecological record itself inherently is scattered. A more detailed, but broadly covering understanding would provide important baseline information for conservation, restoration and rewilding efforts worldwide (*cf.*, e.g., Svenning *et al*., 2016; Fløjgaard *et al*., 2021). Here, a macroecological modelling approach has much to offer. It can help reconstructing maps of different aspects of megafauna assemblage structure (Faurby & Svenning, 2015), and by comparing such estimates with the present faunas allows estimating changes in features relevant to ecosystem functioning, e.g., plant migration rates (Fricke *et al*., 2022b) and trophic network structure (Fricke *et al*., 2022a). In addition, past megafauna distribution reconstructions help forecasting future scenarios of species range shifts in the face of predicted environmental changes and restoration interventions, notably by over-coming anthropogenic truncation of climate niches (Jarvie & Svenning, 2018; Sales *et al*., 2022). While spatial-explicit maps of distributions of both extant and late-Quaternary extinct mammal species with such applications in mind exists (Faurby *et al*., 2020), these maps are limited to relative coarse resolutions and specific, limited time frames, in large part due to limitations of the fossil record.

To overcome these limitations, we here provide detailed estimates of megafauna species ranges for the Last Interglacial (LIG; 129,000-120,000 years ago) in Europe, supported by the region’s extensive literature on megafauna fossils. The LIG is a period of the Pleistocene with climate relatively similar to the present, yet preceding the arrival of *Homo sapiens* in Europe. Hence, it is of interest as a natural baseline for better understanding and managing European nature (Svenning, 2002). The LIG European megafauna community shows continuity with that of previous Pleistocene interglacials, with successful recovery of the same or analogue regional communities following the preceding glacial climate cycles (Nenzén *et al*., 2014; Schreve, 2019). While *Homo neanderthalensis* was widespread in Europe during the LIG, and possibly had local, short-term disturbance on megafauna abundance (Rosell *et al*., 2017; Dembitzer *et al*., 2022), no selective megafauna extinction occurred globally or in Europe prior to the arrival of *Homo sapiens* (Smith *et al*., 2019). Since temperatures were just slightly warmer than at present in the northern Hemisphere (Otto-Bliesner *et al*., 2021), the period has been indicated as the closest in time ecological analogue for the present and near-future Europe, but with an intact megafauna diversity and no widespread human-caused habitat transformations (Svenning, 2002).

Here, we developed detailed estimates of LIG megafauna species distributions in Europe using a fossil-validated hindcasting species distribution modelling approach (Svenning *et al*., 2011), to provide a first spatially-explicit quantification of the megafauna losses in species richness, community biomass (assemblage body mass sum) and functional diversity relative to the present. Furthermore, based on trait-informed species-specific estimations, we test the hypothesis that megafauna losses from the LIG to the present have dramatically reduced the magnitude of potential vegetation and meat consumptions by herbivores (*cf.* Pedersen *et al*., 2020) and carnivores in the European context, likely with strong effects on ecosystem structure and functioning in absence of other human impacts. As the LIG and the present are not climatically identical, we furthermore use a modelling approach to test whether these climatic differences have had a substantial impact on the average levels and patterns of megafauna species richness, community biomass and functional diversity.

## 2. Material and methods

The study area comprises the European mainland west to the Urals (mountains and river) including the Caucasus and Asia Minor (Figure S1). The European mainland geography, particularly coastlines, were considered comparable between the LIG and the present at the resolution of the work due to a relatively similar sea-level (just 5-10 meters above today’s; Dyer *et al*., 2021).

### 2.1 Megafauna distribution data

We retrieved the ‘present-natural’ range of every megafauna (wild terrestrial mammals ≥10 kg) occurring in Europe during the LIG from PHYLACINE v1.2.1 (Faurby *et al*., 2020) (SI Appendix 1 for species selection criteria; Table S1 for the list of 57 species considered at any point in the study). Present-natural ranges are estimates of current-day mammal species distributions, as they would be if *Homo sapiens* disturbance had never occurred. These estimations have been produced globally for all late-Quaternary mammal species at a resolution of 96.5-km grid cell by applying a combination of range adjustments on IUCN historical range maps of still-existing mammals, considering evidence of human-caused extinctions from literature, and co-occurrence based modelling for extinct species (Faurby & Svenning, 2015; Faurby *et al*., 2020).

We collated a record listing latitude, longitude, and name of the excavation site of LIG fossils in Europe (Figure S1; Table S3 and S5) for 38 megafauna species, performing a literature review of studies describing stratigraphic layers associated with the ‘Eemian optimum’ (see section 2.3 for more details). However, we also included few records collated by two studies from eastern Europe, which considered a wider temporal-span for the LIG due to the scarcity of fossils for this region reported in the English-language literature. The review was conducted with Google Scholar using the keywords ‘Eemian’, ‘Last Interglacial’, ‘LIG’, ‘MIS 5e’, ‘Ipswichian’, ‘Mikulino’, in combination with the words ‘fossil(s)’, ‘record(s)’, ‘stratigraphic layer(s)’, ‘(mega)fauna’, ‘mammals’. Each fossil record’s description was carefully evaluated in reliability, crosschecking between references if possible. Moreover, we only included records with geographic coordinates or site’s name available in the referred literature.

We collected the present range for all extant European wild megafauna species (SI Appendix 1 for species selection criteria; Table S1 for the list of 57 species considered at any point in the study) from the International Union for Conservation of Nature (IUCN) Red List of Threatened Species (www.iucn.org; accessed June 2021). We added the species *A. lervia*, *C. nippon*, *H. inermis,* and *O. virginianus* based on Linnell *et al*. (2020), since the European ranges of these introduced species were not reported by the IUCN.

### 2.2 Megafauna functional traits and estimated population densities

First, we classified species as herbivores (Table S6) or carnivores (Table S7) based on the HerbiTraits (Lundgren *et al*., 2021) and the CarniDIET (Middleton *et al*., 2021) databases. From these databases, we also collected functional traits for each species, adding further information on carnivores by Dalerum (2013). Functional traits of *P. spelaea* were not available, but estimated from Sandom *et al*. (2018). We also collected average body mass and percentage of plants and meat in the diet for each species from PHYLACINE v1.2.1 (Faurby *et al*., 2020). We got average species population densities as estimated in conditions of low human impact and ‘field metabolic rate’ for each species from Pedersen *et al*. (2020). Importantly, population densities were assumed constant for the species across their estimated range since quantitative estimation for the whole continent of Europe are missing in the present and are not reliable for the LIG given data scarcity.

### 2.3 Paleoclimate of the LIG and climate of the present

For the LIG paleoclimate, we calculated twelve bioclimatic variables based on climate variables obtained from a set of paleoclimate simulations. These simulations targeted the peak of highest temperature between the two last glacial eras, reached 127,000 years ago (i.e., the ‘Eemian optimum’). They were performed with six climate models (Table S2), each forced according to the protocol for experiment *lig127k* of the Paleoclimate Modelling Intercomparison Project phase 4 (PMIP4; Otto-Bliesner *et al*., 2017). We used the climatological mean of monthly values of the following climate variables: total precipitation, near-surface air temperature, minimum daily near-surface air temperature. We downscaled the climate variables to a resolution of 30-km grid cell. We corrected biases in the climate variables from the models. For this, we compared these variables between the historical simulation with the same climate models and the dataset CHELSA V2.1 (Karger *et al*., 2021), also aggregated at the resolution of 30-km grid cell, as both datasets represent the period 1981-2010. Based on this comparison, we applied a delta correction to the temperature results of the climate model, and a ratio correction to the precipitation results, similar to the procedure applied in, e.g., Scussolini *et al*. (2020). With the bias-corrected variables, we calculated twelve bioclimatic variables (Table S2). LIG paleoclimate is warmer than 20^th^ century average but similar to the early 21^st^ century average climate, thus providing an analogue for global warming thus far (Sánchez Goñi *et al*., 2012), although spatial and seasonal differences in temperature and precipitation were likely quite accentuated (Scussolini *et al*., 2019; Otto-Bliesner *et al*., 2021). For the *present climate*, we retrieved the same bioclimatic variables (Table S2) from CHELSA V2.1 (Karger *et al*., 2021) for the period 2010 – 2020.

### 2.4 Modelling European megafauna species distribution during the LIG

We estimated the distribution of the LIG European megafauna by joint inference of hindcasting species distribution modelling (SDM), fossil records, and geographical constraints on species dispersal.

For the SDM, we generated pseudo-occurrences and pseudo-absences using the species’ present-natural range. We chose this approach since, despite our extensive literature review, the collated fossil record was too scarce to provide reliable estimates of species’ realized niche (Wisz *et al*., 2008). Present-natural ranges can serve as spatially explicit representations of the multi-space environmental niche of a species in modern-like environmental conditions (Jarvie & Svenning, 2018). One pseudo-occurrence was generated per grid cell within the present-natural range at its original resolution (i.e., raster cell of 96.5-km grid cell at 30° latitude) as well without of the present-natural range at the same resolution, limited to a surrounding range buffer with extension equal to one further grid cell. We deleted pseudo-occurrences above 2,000 m following Berti & Svenning (2020), since this elevation represents a spatial constraint for most species that is not accounted for in present-natural ranges. For some species with a very small present-natural range, the number of pseudo-presences generated were too small to use in a SDM. For these, we merged their range with that of a closely related species with similar environmental preferences and treated them as one “species group” (for example grouping *R. rupicapra* and *R. pyrenaica* in a *Rupicapra* spp. group). This concerns 19 species merged into eight groups (see SI Appendix 2 for further details). To estimate the environmental niche for every species, we used the *present climate* predictors retrieved for the decade 2010 – 2020 (Table S2). We used four SDM algorithms to run the models: ‘Bioclim’, ‘Boosted Regression Trees’ (BRTs), ‘General Additive Models’ (GAMs), and Maximum Entropy Modelling (MaxEnt). We run the models using the package *sdm v1.1-8* (Naimi & Araújo, 2016) in R v4.1.0 within and ensemble framework. For each algorithm, the modelling procedure was repeated three times assessing its performance through five-fold cross-validation via computing the area under the curve of receiver operating characteristics (AUC-ROC; see SI Appendix 3 and SI ODMAP protocol - Zurell *et al*., 2020 - for further details). We used the AUC-ROC modelling performance of each single model to weight its contribution to the ensemble prediction, and we computed the mean AUC-ROC as overall evaluation of modelling performance for each species.

The ensemble results for the species were projected in the present climate and in the LIG at a spatial resolution of 30-km grid cell (see SI Appendix 3 for rationale). Amongst the available LIG paleoclimate simulations (Table S2; Otto-Bliesner *et al*., 2021) we utilized the ‘GISS-E2-1-G’ model since it accurately captures the extension of oceanic climate towards the east during the LIG period, as indicated by direct paleobotanical proxies (Pearce *et al*., Highly heterogeneous vegetation characterised the temperate forest biome before *Homo sapiens;* under review). The projected predictions were then converted to binary (presence/absence) outputs (SI Appendix 3 for details). The quality of LIG predictions was evaluated through the fossil record of LIG megafauna (Table S5). We accounted for omission-errors only since the probability of finding fossils reflects environmental conditions favouring fossilization rather than the species most likely distribution (Varela *et al*., 2011). Specifically, we (1) counted the proportion of fossils located inside the species-corresponding range, and we (2) systematically checked whether the model failed in predicting the correspondence of fossils and estimated species range across ten European macro-regions that we defined to encompass the study area (Figure S1; Table S3). The modelling evaluation was performed for 37 species, those for which we both estimated the LIG distribution and retrieved fossil records.

The resulting SDM predictions are estimates of species’ potential – rather than realized – distributions. To improve our estimate of the realized distribution of each species, we cropped the predicted ranges using knowledge on the long-term geographical distribution of each species in the Pleistocene from the literature. We compared the estimated LIG megafauna ranges to the Pleistocene megafauna geographic history as reported in Kurtén (1968), to find geographical constraints in species dispersal throughout the study area. Kurtén (1968) describes fossil assemblages in different areas of Europe, with a focus on western and central regions, over two million years. If such constrains were evident for a species, i.e. its Pleistocene fossils were never found in a particular area, we trimmed the estimated range accordingly (Table S4 for details). Although Kurtén (1968) may have missed evidences of species presence in areas outside the defined boundaries, we considered this source as the most complete to integrate our estimations of potential ranges at ‘high’ resolution with evidences of large scale species endemicity. In addition, we accounted for topography as habitat constraint for some species adapted to steep terrain (SI Appendix 3 for details). Range trimming was not possible for the species not descripted in Kurtén (1968) yet included in our study since, although endemic of Asia, had their range extending in the East European Plain or in Asia Minor, which are parts of our study area (Figure S1).

### 2.5 Quantifying the decline in species richness, community biomass, and functional diversity from the LIG to the present

For both the LIG and the present, we counted the number of megafauna species occurring in each grid cell across the study area, quantifying species richness patterns. For the LIG, we used estimated ranges, while for the present we used ranges collected from the IUCN and Linnell *et al*. (2020) (section 2.4).

For both the LIG and the present, we estimated the biomass occurring per grid cell for each species independently by multiplying average species body size for species population density in ‘natural’ conditions (Table S6 and Table S7 for functional data). We then calculated community biomass patterns summing the biomass of all species in every grid cell across the study area, based on species richness distribution.

For both the LIG and the present, we calculated functional diversity (FD) using the functional traits within species assemblages in each grid cell across the study area based on the species richness distribution (Table S6 and Table S7 for functional data). Considered FD traits and their relative importance (rel. imp.) in FD computation for herbivores, following Schowanek *et al*., (2021), were: i) % of graminoids in diet (0.5 rel. imp.); ii) % of fruits or other plant fractions in diet (0.5 rel. imp.); iii) gut fermentation efficiency (1 rel. imp.); iv) body mass (2 rel. imp.). Considered functional traits and their rel. imp. for carnivores were: i) hunting group size (2 rel. imp.); ii) mean prey size (1 rel. imp.); iii) maximum prey size (1 rel. imp.); iv) body mass (2 rel. imp.); v) selectivity in diet (i.e. assessed spectrum of animal prey considering census by Middleton *et al*., (2021); 1 rel. imp.). FD was calculated as ‘functional richness’ for herbivores and carnivores in a scale from 0.00 (relatively null traits diversity) to 1.00 (relatively full traits diversity). Calculated outputs were then spatially projected across the study area using the *package FD 1.0-12* (Laliberté *et al*., 2014) in r v4.1.0.

The patterns of megafauna richness, biomass and functional diversity were then compared between the LIG and the present by subtracting values of the present from values of the LIG for each grid cell across the study area.

### 2.6 Quantifying the decline in contribution to biogeochemical fluxes (vegetation and meat consumptions) from the LIG to the present

To calculate potential vegetation consumption in kgC*km^-2^yr^-1^ (*PVC*) for each herbivore species independently, we applied the formula in Pedersen *et al*. (2020) for each grid cell in the study area:

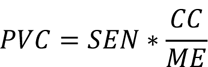

where the species energy needs in kJ*km^-2^yr^-1^ (SEN) is calculated for a period of one year by multiplying estimated species field metabolic rate with estimated population density, and percentage of plants in the diet (data in Table S6; details on the equation in Pedersen *et al*., 2020). *CC* in kgC*kgDM^-1^ is the percentage of carbon contained in dry vegetation matter (*DM*) (45% on average; *SD* = 5.23; Ma *et al*., 2018), and *ME* in kJ*kgDM^-1^ is the metabolic energy available in the selected dry mass of the diet (see Pedersen *et al*., 2020 for further details).

To calculate potential annual meat consumption in kg*km^-2^yr^-1^ (*PMC*) for each carnivore species independently, we used the equation:

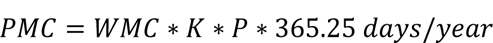

Where wet meat consumption in kg*day^-1^ (*WMC*) is estimated in kilojoules as daily energy intake scaling with body mass, using the pairwise equation in Carbone *et al*. (2007), multiplied by percentage of meat in the diet (*K*) and estimated population density per square kilometre (*P*) (data in Table S7), multiplied by the numbers of days per year. Daily energy intake was then transformed in kilograms (kg) of daily consumed wet meat by applying the caloric conversion for food types presented in SI of Carbone *et al*. (2007) (6,682 kJ/kg for small vertebrate prey and 10,050 kJ for large vertebrate prey).

*PMC* and *PVC* values were summed respectively across all species per grid cell base on species richness distribution, in this case splitting herbivores and carnivores, and compared between the LIG and the present.

### 2.7 Testing megafauna habitat shift due to difference in climate between the LIG and the present

We estimated to what extent the climate difference between the LIG and the present would shift LIG European megafauna ranges and thus the distribution in species richness and community biomass. We did this to test for the potential impact of climate on the average megafauna diversity in Europe from on period to the other. Absence in climate difference would imply a primary role of *Homo sapiens* during the late-Quaternary in creating the actual differences observed. We re-calculated species richness and community biomass patterns as described in sections 2.5, but with megafauna ranges estimated from the projection of LIG megafauna ranges to present-day climate (referred to as ‘present-projected LIG ranges’). We then tested for statistically significant difference in the computed patterns of species richness and community biomass comparing these metrics calculated from 1) ‘present-projected LIG ranges’ *vs* ‘LIG ranges’, and 2) ‘LIG ranges’ *vs* ‘present ranges’. For this, we used the paired Student’s t-test. Importantly, since the test do not distinguish the distribution of the t-test and the normal distribution with more than 30 samples (Kim, 2015), we randomly gathered from the re-computed maps of species richness and community biomass values from the same 30 grid cells in the compared patterns, repeating the process at each t-test iteration.

## 3. Results

### 3.1 The distribution of European megafauna during the LIG

We gathered 365 fossil records for 38 species of European megafauna of the LIG for which we also estimated LIG distribution: the most represented species were *C. elaphus* (n = 35), *C. fiber* (n = 27), *D. dama* (n = 21), *P. antiquus* (n = 20), and *C. capreolus* (n = 19) (Table S3; Table S5). The evaluation of modelling performance to predict LIG megafauna distributions showed an average AUC of 0.82 (± 0.04 Standard Deviation). Omission error evaluation on LIG predicted megafauna ranges using the fossil record showed that 21.7% (n = 79 on a total of n = 365) of the fossils were predicted outside of the correspondent species range. The mean distance of these fossils from the closest point on the modelling estimated range perimeter was only 63.2 km (± 103.1 km SD), however.

### 3.2 Megafauna species richness, community biomass, and FD from the LIG to the present

During the LIG, estimated mean megafauna species richness across Europe was 22.0 species per of 30-km grid cell (95% Inter-quantile Range: 13.0 – 27.0) (Figure 1). In contrast, at present the mean species richness is only 5.5 species per grid cell (95% IR: 2.0 – 9.0) (Figure 1). Mean loss in species richness from the LIG to the present is estimated as 74.3% (± 9.9% SD; Figure 1), with relevant differences amongst present-day countries (Figure S2).

**Figure 1.**
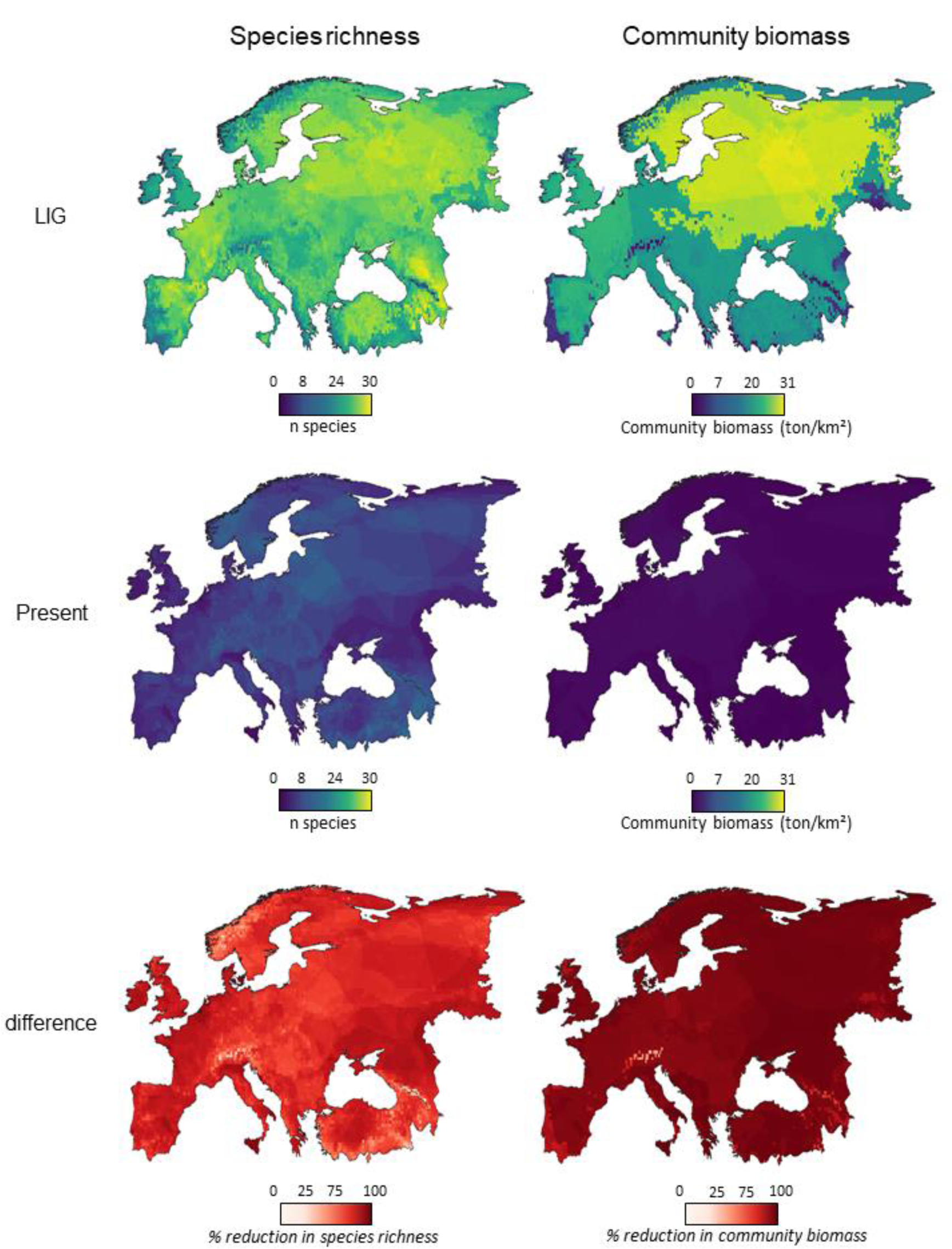
Comparison of megafauna species richness and community biomass between the LIG (cc. 127,000 years ago) and the present (decade 2010 – 2020). LIG species richness was obtained by stacking estimated 48 LIG megafauna ranges (available in SI), while present species richness was obtained by stacking expert-based ranges of the 30 extant European megafauna. From species richness distribution, community biomass was obtained multiplying each species by their body weight and estimated population density in ‘natural conditions’.

During the LIG, estimated mean megafauna community biomass across Europe was 21.7 ton/km^2^ (95% IR: 4.1 – 30.4; Figure 1), with highest biomass (≥25 ton/km^2^) across the East European Plain and Fennoscandia, and lowest biomass (<10 ton/km^2^) along the Mediterranean and Atlantic coasts. In stark contrast, mean biomass in the present is 0.6 ton/km^2^ (95% IR: 0.0 – 1.3; Figure 1), with highest biomass (>1.0 ton/km^2^) in northern parts of east Europe, and lowest biomass (<0.25 ton/km^2^) in the Mediterranean area. Estimated mean loss in megafauna community biomass from the LIG to the present is 96.7% (± 4.1% SD; Figure 1), with dramatic losses everywhere, but least so in mountain ranges.

Mean estimated LIG functional diversity, on a scale from 0.00 to 1.00, was of 0.64 (95% IR: 0.36 – 0.79; Figure S3) for herbivores and of 0.65 (95% IR: 0.16 – 1.0; Figure S3) for carnivores. In the present, mean FD is only 0.05 (95% IR: 0.0 – 0.15; Figure S3) for herbivores and 0.17 (95% IR: 0.0 – 0.48; Figure S3) for carnivores. Overall, Europe has lost a grid cell mean of 59.1% (± 11.8% SD) of the large-herbivore FD and 48.2% (± 25.0% SD) of the large-carnivore FD from the LIG to the present (Figure 3).

**Figure 2.**
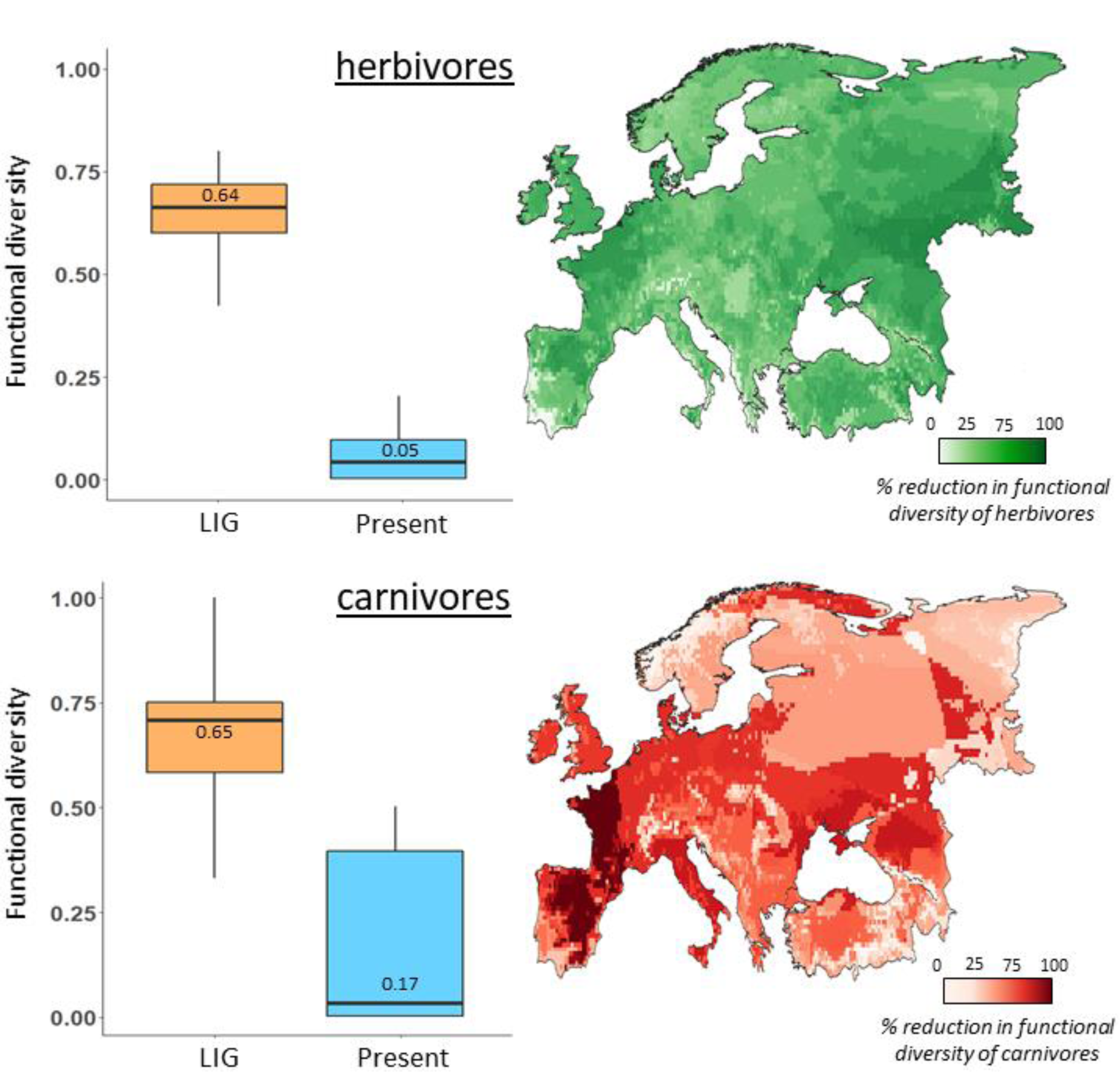
Comparison of functional diversity (FD) between the LIG and the present, considering 37 herbivore species (Table S6) and 14 carnivore species (Table S7) occurring in one or in both periods (NB. *U. arctos* was considered as both herbivore and carnivore). Boxplots represent the variability in functional diversity across the study area. The maps show the estimated relative loss in FD between the LIG (cc. 127,000 years ago) and the present.

**Figure 3.**
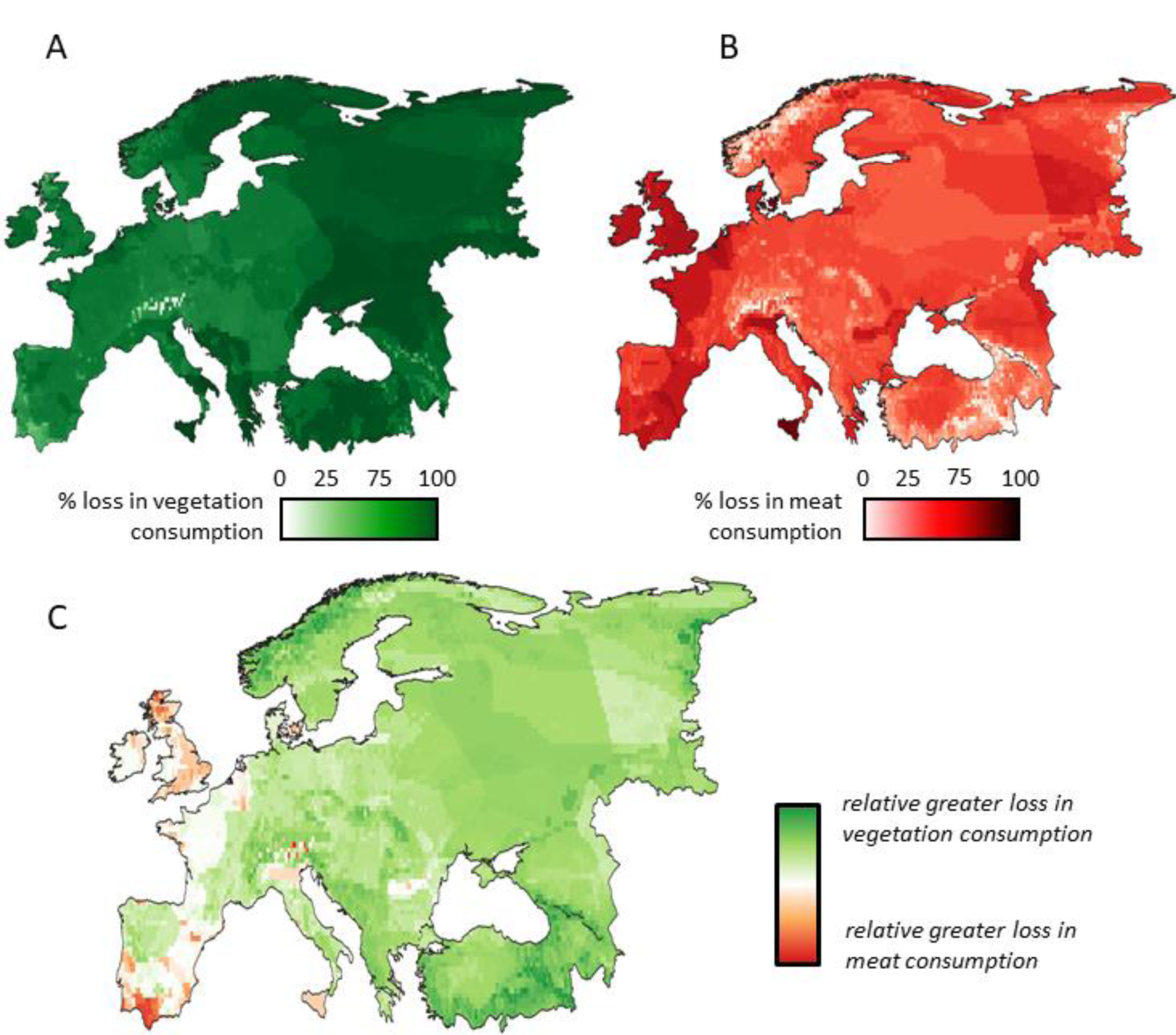
(A) Estimated loss in dry vegetation consumption and in (B) wet meat consumption from the LIG to the present. (C) Relative greater loss in either vegetation or meat consumption, i.e., whether a disproportional demise in herbivores or carnivores consumption happened in a particular area based on the consumption demise ratio. This was calculated as (% loss in vegetation consumption - % loss in meat consumption per unit area); resulting positive values represent greater loss in vegetation consumption, and resulting negative values represent greater loss in meat consumption, on a scale −100 to 100).

**Table 1.** Summary of the megafauna biodiversity metrics calculated, with estimated results for the LIG and the present, and quantified loss between the two periods.

| Megafauna biodiversity metric | LIG <sup>A</sup> | Present <sup>A</sup> | Loss |
| --- | --- | --- | --- |
| <u>Species richness</u><sup>1</sup> | 22.0 (13.0 – 27.0) | 5.5 (2.0 – 9.0) | 74.3% |
| <u>Megafauna community biomass</u><sup>2</sup> | 21.7 (4.1 – 30.4) | 0.6 (0.0 – 1.3) | 96.7% |
| <u>Functional diversity (FD) herbivores</u><sup>3</sup> | 0.64 (0.36 – 0.79) | 0.05 (0.0 – 0.15) | 59.1% |
| <u>Functional diversity (FD) carnivores</u><sup>3</sup> | 0.65 (0.16 – 1.0) | 0.17 (0.0 – 0.48) | 48.2% |
| <u>Vegetation consumption</u><sup>4</sup> | 31.1 (10.4 – 42.6) | 2.5 (0.4 – 5.8) | 91.1% |
| <u>Meat consumption</u><sup>4</sup> | 1.0 (0.5 – 1.3) | 0.1 (0.1 – 0.6) | 61.2% |

A - Results reported as mean with 95% interquartile in brackets. 1 - Computed as number of species. 2 - Computed as ton/km<sup>2</sup>. 3 - Computed as index from 0 (null traits diversity) to 1 (full traits diversity). 4 - Computed as ton/km<sup>2</sup>/year of dry vegetation matter and wet meat.

### 3.3 Vegetation and meat consumptions by megafauna from the LIG to the present

During the LIG, potential mean annual vegetation consumption – as estimated from the species assemblage and their traits – was 31.1 ton/km^2^/year (95% IR: 10.4 – 42.6). In the present, potential average vegetation consumption is only 2.5 ton/km^2^/year (95% IR: 0.4 – 5.8). We estimated that potential average vegetation consumption in Europe is diminished by 91.1% (± 7.4% SD) from the LIG to the present, with biggest differences (≥90%) in the east and the smallest differences (≤50%) are found in mountainous regions (Figure 3-A).

During the LIG, estimated potential mean meat consumption across Europe was of 1.0 ton/km^2^/year (95% IR: 0.5 – 1.3), while current potential mean meat consumption is only 0.1 ton/km^2^/year (95% IR: 0.1 – 0.6) (Figure S4-B). In terms of relative loss, estimated potential meat consumption is diminished by 61.2% (± 17.2% SD) across Europe, with the biggest differences in Western Europe and the smallest differences in mountains (Fig. 3-B).

**3.4 Estimated megafauna habitat shift due to different climate between the LIG and the present** To estimate the effects of overall faunal differences and removing any effects of climate differences between the LIG and the present, we also compared species richness and community biomass patterns constructed by (1) ‘LIG ranges’, (2) ‘present-projected LIG ranges’, and (3) ‘present ranges’ (Figure S5). On mean, megafauna species richness computed with present-projected LIG ranges is 0.7 species/grid-cell higher than megafauna species richness computed with LIG ranges, without estimated statistical difference (t(29) = −1.34, p-value = 0.19). Megafauna species richness computed with LIG ranges is 16.5 species/grid-cell higher than megafauna species richness computed with present ranges, with estimated high statistical difference (t(29) = 39.67, p-value = <0.001). On mean, megafauna community biomass computed with present-projected LIG ranges is 0.035 ton/km^2^ higher than megafauna community biomass computed with LIG ranges, without estimated statistical difference (t(29) = −0.05, p-value = 0.96). Megafauna community biomass computed with LIG ranges is 21.1 ton/km^2^ higher than megafauna community biomass computed with present ranges, with estimated high statistical difference (t(29) = 30.31, p-value = <0.001).

So, there is weak and non-significant difference in the potential mean species richness and community biomass patterns driven solely as the result of LIG vs. present-day climate differences, while in reality there are very strong differences in the megafauna ranges and thus in the diversity of ecological effects patterns between the two periods, as assessed in this study.

## 4. Discussion

Wild megafauna diversity and associated potential ecological effects in the ecosystems of Europe are dramatically reduced in the present day compared to the Last Interglacial (LIG). Megafauna losses have occurred everywhere across Europe, with relatively small geographic variability, with the least losses in mountainous regions. Species richness is on average 74% lower and megafauna community biomass is 90% lower everywhere. Functional diversity loss for both herbivores and carnivores exceeds 50% across most of Europe. Importantly, the associated ecosystem impacts as indicated by our estimates of potential vegetation and meat consumptions are similarly reduced, by 91% and 61% on average across Europe. These massive reductions in megafauna diversity and potential functional effects mean that contemporary natural ecosystems in Europe strongly deviate from the evolutionary norm given the uniqueness of faunal downsizing in the window of time between the LIG and the present (Smith *et al*., 2018). Importantly, our results also show that the small climate differences between the LIG and the present included in our modelling do not explain these reductions, and would instead have led to small increase in megafauna richness in the present, in the absence of the human-linked late-Quaternary extinctions (Sandom *et al*., 2014a; Smith *et al*., 2019)

### 4.1 Loss in ecological functions sustained by megafauna herbivores

The potential effects of megafauna herbivores on European ecosystems have reduced dramatically from the LIG to the present. Notably, we found that the functional diversity of herbivores in Europe is reduced by 59% on average, and overall potential vegetation consumption in wild ecosystems is 91% lower. By comparing the estimated diversity and biomass of the LIG European megafauna herbivores guild with data from today’s sub-Saharan African reserves (Hempson *et al*., 2015; Fløjgaard *et al*., 2021), we found fairly similar or even higher values, likely leading to comparable processes of primary-consumers control on vegetation succession and fire prevalence. As in other parts of the world (Schowanek *et al*., 2021), late-Quaternary extinctions therefore strongly reduced megafauna herbivore assemblages in Europe until the present, and this trend is not recent (Smith *et al*., 2019). This is in line with the analyses on British dung beetle assemblages from the LIG, which indicated more frequent high abundance of large herbivores than during the early Holocene (Sandom *et al*., 2014b) and strong down-sizing of dung-beetle communities subsequently (Schweiger & Svenning, 2018). Such high abundance of large herbivores have potential to generate heterogeneous vegetation including open and semi-open components (Bakker *et al*., 2016), as also seen on fertile soils in European rewilding areas today (Cornelissen *et al*., 2014). Temperate Europe indeed shows evidence of substantial presence of open vegetation during late Pleistocene interglacials, including the LIG, especially in floodplain areas, on marginal soils and dry climates (Svenning, 2002; Sandom *et al*., 2014b). High heterogeneity in vegetation is generally associated to high species richness (Vera *et al*., 2006). Many European species depend on open and semi-open vegetation, e.g., three groups of light-demanding plants: many forb species, many thorny shrubs, and tree species that cannot regenerate in present-like deep shade, such as oaks (*Quercus* spp.) and hazel (*Corylus avellana*) (Pykälä *et al*., 2005; Bobiec *et al*., 2018).

### 4.2 Loss in ecological functions sustained by megafauna carnivores

The functional diversity of carnivores in Europe is reduced by 48% on average, and potential overall meat consumption is 61% lower in the present compared to the LIG. While these declines are not as steep as for herbivores, the demise of carnivores in Europe compared to an evolutionary baseline likely have had nontrivial consequences for ecosystems. Meat consumption was particularly diversified in the LIG and abundant in some areas of carnivores co-occurrence, with clear niche partitioning thus competition avoidance (Konidaris, 2022) due to heterogeneity in body weights, diet, and social structure. Cave lion (*P. spelaea*), whose ecological niche was similar to today’s African lions but was physically much bigger (Manuel *et al*., 2020), was widespread in Europe. The diet of this animal was oriented toward large ungulates such as equids (*Equus* spp.) and aurochs (*B. primigenius*), but also included the cave bear (*U. spelaeus*) (Bocherens *et al*., 2011). Other top-carnivores in Europe during the LIG were the spotted hyena (*C. crocuta*) and the grey wolf (*C. lupus*), pack hunters feeding on mid-to-high size herbivores, including scavenging on bones. We estimated these species as widely distributed across Europe, also in accordance with the fossil record, most likely hunting in wide-open areas (Diedrich, 2014). The diversified LIG guild of large carnivores was also composed of ambush predators such as the Eurasian lynx (*L. lynx*) and the leopard (*P. pardus*). The presence of ambushing carnivores triggers fear-driven mesoherbivore aggregations in open areas and thus redistribution of soil fertilization by faeces (le Roux *et al*., 2018). Interesting, most of these carnivores co-occurred in Western Europe where at present they are all virtually absent, determining a dramatic functional diversity drop in this region. Particularly in British Isles, this has also been associated with the overabundance of herbivores such as deer, which overgrazing causes homogenization of the landscape, but also lead to conflict with humans relative to the danger of car collisions, crop damage, and potential for the spread of diseases (Côté *et al*., 2004).

At the present, fundamental ecological functions provided by extant carnivores are largely relegated to remote areas and mountainous regions, such as in the central Apennines, the Carpathians, the north of Scandinavia, but also in lowlands between Belarus and Poland. In these areas, carnivores still exert density-mediated and behaviourally-mediated effects of their prey, with cascading consequences on the lower trophic levels (Kuijper *et al*., 2013). Only the Balkans, Turkey and Caucasus have conserved levels of meat consumption and carnivores’ functional diversity that approach the LIG. However, local people frequently poach carnivores in the southeast of Europe given high intolerance toward coexistence in shared landscapes (Ripple *et al*., 2014; Ghoddousi *et al*., 2020). Yet, large carnivores have been a focus for conservation efforts during the last decades, and are generally in a positive trend of comeback particularly in south and central Europe (Chapron *et al*., 2014).

### 4.3 Reliability of estimated LIG European megafauna diversity

The record of LIG megafauna fossils is scarce for most species. A more exhaustive record would allow the combination of habitat models trained directly with LIG environmental features and of co-occurrence analyses to infer spatial competition. We were unable to reliably perform such analysis for most of the species given the scarcity of data, particularly for eastern Europe. Investigating literature in non-English languages would likely alleviate this problem to some extent, but strong under-sampling and geographic bias is likely to be a persistent feature of the fossil record. Notably, some of the species have a very sparse fossil record, a situation that is unlikely to be remedied anytime soon.

While the number of omission errors, i.e., fossils out of estimated species ranges, were relatively high, the average distance of these fossils from the estimated range perimeter was <100 km, i.e., well within the range of the movement distances of individuals of most megafauna species. Hence, effectively our range estimates did not result in substantial omissions in most cases. There was, however, a LIG fossil record with substantial omission errors, namely *Mammuthus primigenius* (830 km from the perimeter of the predicted range). This record was reported by Markova (2000), which defines the LIG broadly from 140,000 to 120,000 years ago. Hence, the fossil could reflect species occurrence from outside the optimal phase of the LIG, but other possibilities such as underestimation of the niche of woolly mammoth (*M. primigenius*) is possible. Importantly, while this species is often understood as a cold-climate associated species it has LIG records from southern parts of the East European Plain as do woolly rhinoceros (*C. antiquitatis*) (Markova, 2000) as well as other records from relatively mild climates (West, 1969; Álvarez-Lao & Garcia, 2012).

Due to lacking reliable data on LIG population densities for individual megafauna species or their species variation, we had to estimate these based on their traits (Pedersen *et al*., 2020) as constant population density values throughout the species ranges. Range-wide constant population densities are unlikely in most cases (Martínez-Meyer *et al*., 2013), but should here be seen simply as a general indication of the species’ typical potential density. Similarly, the dependent estimated consumption rates should therefore also only be seen as generalized estimates of each species’ potential effect.

### 4.5 Key-message for restoration ecology and conclusion

With the severe reduction in megafauna diversity and associated potential functional effects from the LIG to the present, a unique phenomenon in the last >10 million years, current European natural ecosystems deviate strongly from their long-term evolutionary conditions. While Europe has experienced a remarkable comeback of megafauna during the last decades (Deinet *et al*., 2013), our results show that present megafauna diversity is still just a small fraction of what has characterized European ecosystems prior to the *Homo sapiens-*linked fauna losses of the last 50,000 years. This faunal simplification has strong implications for European nature, not least in relation to the widespread occurrence of land abandonment and associated passive rewilding (Navarro & Pereira, 2015). Woody densification is a widespread phenomenon in European nature and a threat to a large proportion of Europe’s biota, and is in large part associated with reduced presence of large herbivores in the landscape (e.g., Buitenwerf *et al*., 2018). At the same time, natural areas in some regions experience biodiversity losses linked to extremely high, uniform presence of deer, likely in large part linked to reduced or absent large-carnivore assemblages. These dynamics seem obviously linked to the downsizing and simplification of the European fauna quantified here. At least partially restoring faunal functionality through trophic rewilding interventions (Svenning *et al*., 2016) should therefore be amongst the priorities for the agenda of European countries to safeguard and restore the continent’s biodiversity. A rising number of real-world implementations of trophic rewilding provide empirical support for positive effects on biodiversity, in Europe (e.g., Dvorsky *et al*., 2022) as well as on other continents (Guyton *et al*., 2020; Ratajczak *et al*., 2022). Furthermore, megafauna-based trophic rewilding can also be seen as a contribution to nature-based solutions to climate change via assisting ecological adaptation to climate change and hence maintenance of climate change mitigation contributions, such as vegetation and soil carbon sinks (Malhi *et al*., 2022). Our study thus supports the restoration of megafauna diversity and ecological effects to European natural and semi-natural landscapes as an important countermove against the current environmental crisis.

## Acknowledgments

We thank Emilio Berti and Scott Jarvie for useful early discussions on the study. We also thank Erick Lundgren for his assistance with the analysis of functional diversity and Dereck Corcoran for assistance in processing paleoclimate data.

The work was supported by the project TERRANOVA, the European Landscape Learning Initiative, which has received funding from the European Union’s Horizon 2020 research and innovation programme under the Marie Sklodowska-Curie grant agreement no. 813904. The output reflects the views only of the authors, and the European Union cannot be held responsible for any use which may be made of the information contained therein.

J.C.S. and R.P. considers this work a contribution to Center for Ecological Dynamics in a Novel Biosphere (ECONOVO), funded by Danish National Research Foundation (grant DNRF173) and his VILLUM Investigator project “Biodiversity Dynamics in a Changing World” funded by VILLUM FONDEN (grant 16549) and his Independent Research Fund Denmark | Natural Sciences project MegaComplexity (grant 0135-00225B).

D.N.K. acknowledges funding from: The WSL internal grant exCHELSA, and ClimEx, and the 2019–2020 BiodivERsA joint call for research proposals, under the BiodivClim ERA-Net COFUND program, with the funding organisations Swiss National Science Foundation SNF (project: FeedBaCks, 193907).

## Author statement

M.D. and J.C.S. conceived the ideas. M.D., S.M., S.N. and J.C.S. designed the study. P.S., D.N.K. and R.P. provided key datasets. M.D. implemented the analyses. M.D., S.M. and J.C.S. led the writing. All co-authors contributed critically to results interpretation and manuscript draft.

## Data availability statement

Table S5, Table S6, Table S7, and the estimated LIG ranges of European megafauna are openly available in figshare at https://doi.org/10.6084/m9.figshare.22350907

## Biosketch

**Marco Davoli** is a postdoctoral researcher at Roma La Sapienza University, Italy, who recently obtained his PhD from Aarhus University, Denmark. His research primarily focuses on historical ecology, human-wildlife coexistence, ecological restoration, and megafauna rewilding, with a specific emphasis on European ecosystems. Presently, his work involves establishing favourable reference values of conservation for non-mammalian species within the European Union.

