## Appendix 1 for "Megafauna diversity and functional declines in Europe from the Last Interglacial (LIG) to the present"

### **Appendix 1. Criteria to select European megafauna species of the Last Interglacial (LIG; cc. 127,000 years ago) and the present**

We defined megafauna as wild, terrestrial, non-flying mammals with body weight ≥ 10 Kg (following Sandom *et al.*, 2014a). For the LIG, we considered the 38 megafauna species for which we gathered LIG fossils or listed into Table 15 ´Eemian European mammal fauna´ within Kurtén (1968). This book provides a full description of the fossils record found across western and central Europe and attributed to the Pleistocene, including the LIG (reported as MIS 5e^A^). However, we included in the analysis further 13 species which presence in Europe during the LIG was suspected in Anatolia and East European Plain based on the present natural ranges of PHYLACINE v1.2.1 (Faurby *et al.*, 2020). Notably, if the present natural range of a species overlapped with the study area, we tested for the occurrence of its suitable habitat in Europe under LIG climate. If the model returned indeed suitable habitat we considered the species for the study, otherwise the species was discarded. This gave 51 species potentially occurring in Europe during the LIG. From this total, despite acknowledging its occurrence in Europe during the LIG, we excluded *Bubalus murrensis* from the analyses of ecological effects given the impossibility to estimate its LIG range due to missing present-natural range (Section 1.2.4 main text for rationale about the use of present-natural ranges in the modelling framework). We also excluded *Equus ovodovi* and *Homotherium latidens* since their occurrence in Europe during the LIG is questionable, respectively given phylogeny uncertainty and lacking fossil evidences also in nearby regions. Eventually, we therefore considered 48 LIG megafauna species in the analyses of ecological effects.

For the present, we listed 26 species occurring as wildlife in the study area based on International Union for the Conservation of Nature (IUCN) Red List extant ranges ([www.iucn.org](http://www.iucn.org); accessed June 2021). We considered all megafauna of the present hence including both native and introduced species. We also considered further four species based on Linnell *et al.* (2020), for 30 megafauna for which we collated the present range for the analyses.

For both periods, we excluded the *Homo* genus including both *Homo neanderthalensis and Homo sapiens,* in order to focus the study purely on the species typically considered as megafauna.
