## Appendix 2 for "Megafauna diversity and functional declines in Europe from the Last Interglacial (LIG) to the present"

### **Appendix 2. Further details on the preparation of pseudo-occurrences and pseudo-absences**

We considered eight groups of sister species as belonging to the same taxa, given (A) the small extension of the species present-natural range leading to few occurrence records generated, thus low modelling calibration statistical power; (B) comparable functional traits thus ecological effects; (C) the need to avoid estimated co-occurrence of multiple species that likely occupied the same ecological niche. The eight groups were (1) *Capra ibex* / *Capra pyrenaica* (considered as ‘Alpine *Capra* spp.*’*), (2) *Capra caucasica* / *Capra aegagrus* / *Capra cylindricornis* (‘Caucasian *Capra* spp.*’*), (3) *Capreolus capreolus* / *Capreolus pygargus* (“*Capreolus* spp.”), (4) *Cervus elaphus* / *Cervus canadensis* (‘*Cervus* spp.*’*), (5) *Hystrix indica* / *Hystrix cristata* / *Hystrix refossa (‘Hystrix* spp.*’*; as distinguished from *Hystrix brachyura spp. vinogradovi* that was present in Europe during the LIG, but is not considered in our study given body weight < 10Kg), (6) *Rupicapra rupicapra* / *Rupicapra pyrenaica* (‘*Rupicapra* spp.*’*)*.* Moreover, based on recent literature, we considered (7) *Equus hydruntinus* as a subspecies of *Equus hemionus* (i.e. '*Equus hemionus hydruntinus'*; Boulbes & Van Asperen, 2019), despite these species are distinguished in Phylacine v1.2.1. Furthermore, though acknowledging the possible existence of two distinct *Bison* lineages in the region in the late Quaternary and thus in the LIG (*B. schoetensacki, B. bonasus, B. priscus*), we represented this group (7) as the taxon ‘*Bison* spp.*’*. This to estimate the total range for the group into the explained framework iterated, as Phylacine v1.2.1 merges these species under *B. bonasus* accounting for still unclear, complex phylogenetic relationships within the genus (e.g., *B. schoetensacki* is likely a chronospecies of *B. bonasus;* Palacio *et al.*, 2017)*.*
