## Appendix 3 for "Megafauna diversity and functional declines in Europe from the Last Interglacial (LIG) to the present"

### **Appendix 3. Further details on the modelling of megafauna distribution during the LIG**

We performed hindcasting SDMs by applying a multi-modelling ensemble approach using the R package *sdm v1.1-8* (Naimi & Araújo, 2016) and the following algorithms: Bioclim, Boosted Regression Trees (BRTs), General Additive Models (GAMs), Maximum Entropy modelling (MaxEnt) (Naimi & Araújo, 2016)*.* These widely used presence-absence, presence-background, and presence-only (profile) methods implement SDM through different computations, and perform with different quality based on the nature of the input data and the processed number of records relative to predictors variability. However, they generally outperforms other algorithms of the same type (e.g., Elith *et al.*, 2006; Li *et al.*, 2017). An ensemble approach increases overall modelling performance by building on consensual predictions of habitat suitability amongst different techniques, compensating inherent biases in single algorithms (Grenouillet *et al.*, 2011; Qiao *et al.*, 2015). The contribution of each algorithm to the model was calibrated through evaluating the modelling by five-fold cross partitioning of the training data. In results, we report the evaluation on training modelling considering the mean of evaluation scores among all algorithms for each species.

Before modelling, we tested for multi-collinearity the bioclimatic predictors used in model calibration by calculating the Variable Inflation Factor (VIF). We discarded predictors with a VIF ≥3 (James *et al.*, 2013). Predicted habitat suitability values as probabilistic output were eventually spatially-projected across the study area at a resolution of 30-km grid cell (length of the side of a 900 km² area at 30° latitude; ´Behrmann` projection) using the LIG paleoclimate predictors. We selected a 30km resolution after conducting sensitivity analysis of tested projections at higher spatial resolutions (i.e., 10km, 20km), for which were found oddities resulting from extreme climatic conditions in mountainous areas, not addressable by our course-scale modelling approach. We eventually converted the averaged species estimated ranges projection as presence/absence (binary) using the minimum value amongst the five threshold methods highlighted as best performing by Liu *et al.* (2005), to remain conservative on distributions. These were prevalence, average probability/suitability, sensitivity-specificity sum maximization, sensitivity-specificity equality, and the ROC-plot based approach.

In range trimming (Table S4), we considered the dependence of caprine species on roughed terrain as adaptation against predation, thus their predicted ranges were limited where Terrain Roughness Index (TRI) was ≥ 240 unit (intermediate-to-high terrain roughness; Riley *et al.*, 1999). We considered this as the least roughness required by caprine for suitable habitat based on Acevedo *et al.* (2007). Average TRI at 30 x 30km was calculated using the downscaled 90 x 90m resolution World digital elevation model available from Robinson *et al.* (2014).
